# Monolithic Shape-Shifting Absorbable Implants for Long-Term Contraception

**DOI:** 10.1101/2025.05.18.654764

**Authors:** Jason Li, Benjamin G. Clark, Parmiss Khosravi, Colin Cotter, Jia Y. Liang, Susan R. Ling, Yuyan Su, Kent J. Qi, Flavia Codreanu, Ethan D’Orio, Ivy Tianqin Tan, Julia E. Green, Katarzyna Murawska, Aaron Lopes, Benjamin N. Muller, Alison M. Hayward, Niora Fabian, Andrew Pettinari, Kailyn Schmidt, Benedict Laidlaw, Maria Platero, Ashley Guevara, Joshua D. Bernstock, Stephanie Y. Owyang, Peter R. Chai, Robert Langer, Giovanni Traverso

**Author notes:** Corresponding authors (G.T.), (J.L.).

## Abstract

Reversible contraceptives empower women to prevent unintended pregnancies and enable family planning. However, the need for frequent dosing with pills or injections often leads to suboptimal medication adherence and reduced effectiveness–an issue common to many chronic conditions. Long-acting drug delivery implants offer a compelling alternative by enabling autonomous, multi-year drug release, thereby improving real-world adherence and treatment outcomes. However, user acceptability and access are limited by need for invasive insertion and surgical end-of-life removal, particularly in low-resource settings, as well as by limited drug loading and suboptimal drug utilization efficiency, which constrain both the duration of therapy and the range of drugs that can be effectively delivered.

To address these limitations, we developed the Monolithic Shape-shifting Absorbable Implants for Chronic Care (MoSAIC) platform–a minimally invasive, fully bioresorbable system that integrates compacted drug formulations with a space-efficient device architecture. This approach reduces implant size, eliminates the need for surgical removal, and prolongs therapeutic duration compared to existing implants. We develop compacted formulations of the contraceptive drug levonorgestrel (LNG), and other poorly water-solubility drugs, demonstrating exceptional drug loading (100% w/w) and multi-year sustained drug release *via* surface-mediated dissolution in rats. When incorporated into MoSAIC devices, these formulations enable high-efficiency drug loading and zero-order drug release kinetics with geometrically tunable rates and durations. As a result, MoSAIC systems can be designed to be smaller, less invasive, and/or longer lasting than current contraceptive implants such as Jadelle® and Nexplanon®.

The MoSAIC platform expands access to reversible contraception and supports long-term medication adherence, with the potential to improve health outcomes and quality of life. More broadly, it provides a flexible approach for delivering other potent, low-solubility therapeutics and lays the foundation for a “dose it and forget it” paradigm in chronic disease management, where adherence is designed into the therapy itself.

## Main

Despite significant advances in pharmaceutical development that have transformed the management of chronic health conditions, medication non-adherence remains a major barrier to realizing the full benefits of these therapies. For instance, while reversible contraceptives enable women to prevent unintended pregnancies^1,2^, the necessity for repeated dose administration over extended periods contributes to low adherence and high discontinuation rates^3^. Among college women in the United States, adherence to daily oral contraceptive pills is estimated at only ∼52%, resulting in a drop in effectiveness from 99.7% under perfect use conditions to 91% in typical use conditions^4–6^. Similarly, low adherence rates are observed for a range of other chronic drug therapies including for hyperlipidemia, diabetes, obesity, hypertension, asthma, depression, epilepsy, human immunodeficiency virus (HIV), and tuberculosis^3,7–12^. As a result, suboptimal medication adherence severely compromises treatment effectiveness, leading to high discontinuation rates, poorer health outcomes, and increased healthcare costs^3,4^.

Medication adherence can be improved by developing and adopting new treatment regimens that require fewer and less frequent dose administration events^13^. As a result, extended-release drug delivery systems with progressively longer dosing intervals have been the focus of intense development over the past two decades. Injectable polymer microspheres, crystalline suspension drug depots, oil-based formulations, and *in situ* forming polymer implants are promising extended-release technologies; however, the therapeutic duration achievable with these systems is largely limited to 1-6 months^14,15^. Furthermore, these systems are generally non-retrievable, precluding the possibility of discontinuing therapy if patients develop adverse reactions or change their family planning goals.

Implantable drug delivery devices also offer extended drug release profiles, with the added benefit of being easily retrievable and providing substantially longer therapeutic coverage compared to injectable formulations^16^. For instance, the Nexplanon^®^ and Jadelle^®^ subdermal implants can provide 3 or 5 years of contraception, respectively, and are the most effective form of reversible contraceptives available today^17^. A prospective cohort study showed that 75% of women in the St. Louis area (USA) preferred long-acting reversible contraceptive systems such as implants and intrauterine devices over shorter-acting self-administrable systems (i.e., daily oral pills, long-acting injectables, vaginal rings, patches, and diaphragms) when the barriers of cost, knowledge and access were removed^18^. Thus, long-acting drug delivery implants can offer unique advantages for the management of chronic conditions by leveraging their ultra-long therapeutic durations to shift the treatment paradigm from frequent patient self-administration, which is prone to low adherence rates, to a dose-it-and-forget it strategy where near-perfect adherence is engineered into the system (**Fig. 1A**).

**Figure 1.**
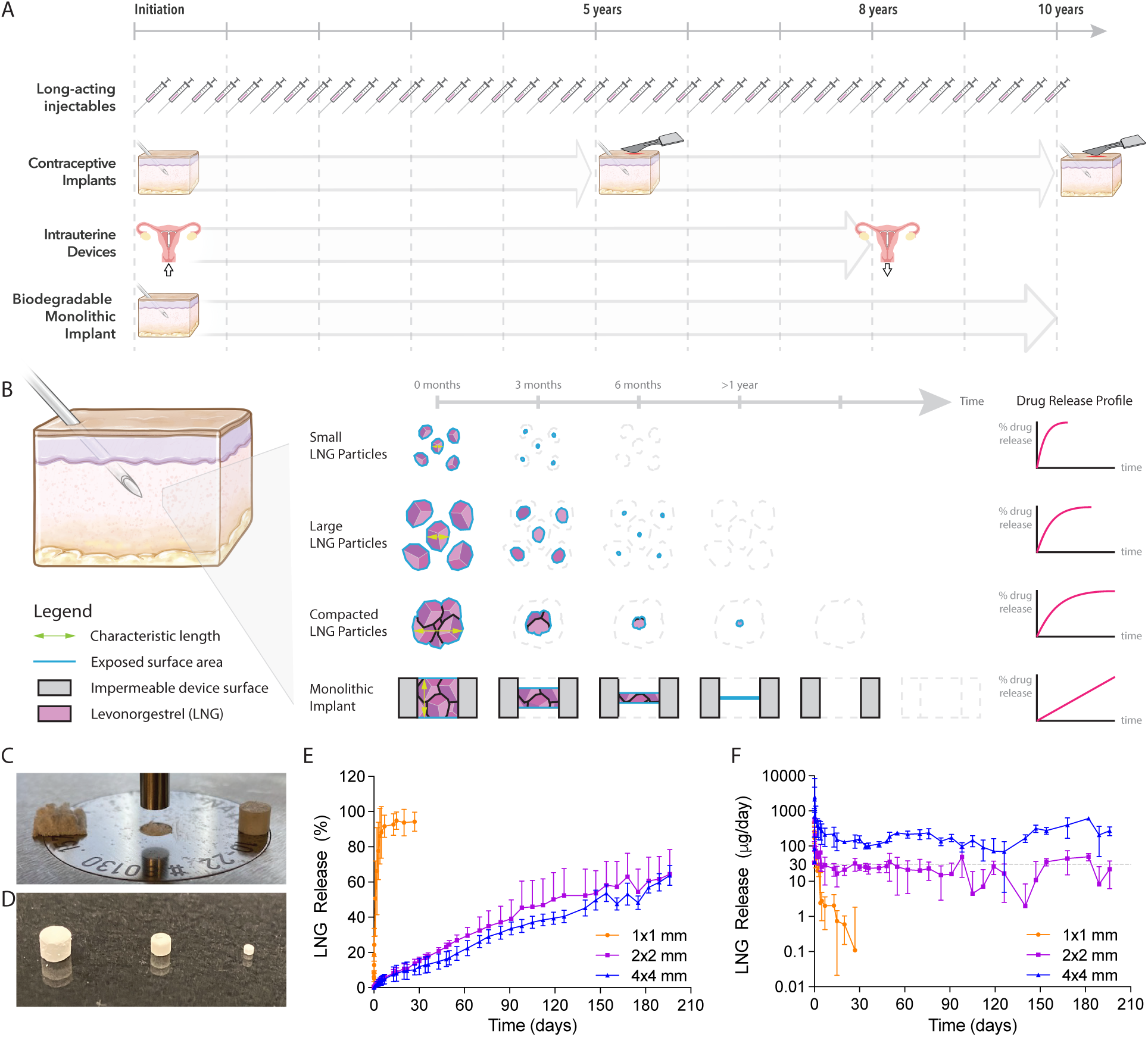
A) Schematic illustrating the dosing regimen and therapeutic duration of current reversible long-acting contraceptive systems compared to the proposed bioresorbable MoSAIC system. B) Schematic illustrating the effect of size and geometry of four different LNG formulations on the drug release profile and therapeutic duration. C) Photograph showing the tablet compression of lyophilized LNG to form compacted LNG formulations. D) Photograph of compacted LNG formulations of three different sizes: 1×1 mm, 2×2 mm, and 4×4 mm. E) *In vitro* drug release profile for the three different compacted LNG formulations in pH 7.4 PBS supplemented with 3% SDS at 37°C under sink conditions and constant agitation. Data plotted as mean ± SD (n≥4). F) *In vitro* drug release rate for the three different compacted LNG formulations. Data plotted as mean ± SD (n≥4).

Despite the improved adherence, effectiveness, and convenience offered by long-acting drug delivery implants, patient adoption and access to these systems can be limited by the need for subcutaneous insertion into the arm using a trocar, and end-of-life surgical removal (**Fig. 1A**)^16,19^. These procedures can be painful, increase risk of infection, and require access to a clinic^19^. Furthermore, the limited volume available for drug loading within these minimally invasive devices drastically restricts the type of drugs that can be delivered, the list of clinical indications that can be targeted, and the achievable therapeutic durations^16^. To address these challenges, we report the development of the **Mo**nolithic **S**hape-shifting **A**bsorbable **I**mplants for **C**hronic **C**are (**MoSAIC**) platform – a minimally invasive and fully biodegradable implant architecture that maximizes the efficiency of drug loading, drug utilization, and drug release of low solubility drugs to enable smaller, less invasive, and/or longer-lasting implants. We focus on the development of MoSAIC for the delivery of levonorgestrel (LNG), a potent progesterone used for long-term contraception, but also demonstrate its applicability towards other chronic conditions which face analogous long-term adherence challenges and technical limitations preventing improved long-term treatment options.

## Results

### Compacted levonorgestrel formulations exhibit sustained drug release kinetics

The MoSAIC platform employs millimeter-sized, compacted formulations of potent, poorly water-soluble drugs to achieve higher drug loading capacities than polymer-drug matrix systems (e.g., Jadelle®, Nexplanon®) and to enable prolonged therapeutic durations compared to injectable microcrystalline suspension drug depots (e.g., Depo Povera®, Sayana Press®, Apretude®). This approach draws inspiration from injectable aqueous crystalline suspension depot formulations, which provide extended drug release through the gradual partitioning of drug molecules from the surfaces of micron-sized particles, with release duration proportional to particle size (**Fig. 1B**)^14,15^. For instance, Depot-Provera CI® is formulated as an aqueous suspension containing medroxyprogesterone acetate particles approximately 11 µm in diameter, which dissolve slowly following intramuscular injection to provide ∼3 months of sustained drug release^20^. While extending drug release beyond this 3-month window by increasing particle size is a potential strategy, manufacturing large, millimeter-scale crystalline drug particles for multi-year efficacy is impractical. We therefore hypothesize that a millimeter-scale monolithic formulation, created by mechanically compacting micron-sized drug crystals, could enable surface-mediated dissolution similar to that of individual particles, thereby facilitating multi-year drug release durations (**Fig. 1B**).

Cylindrical compacted monolithic LNG formulations with diameters of 1 mm, 2 mm, and 4 mm and heights of 1 mm, 2 mm, and 4 mm, respectively, were fabricated by tablet compression (**Fig. 1C, D, Table S1**). The tablets were composed entirely of drug (i.e., 100% wt.) and had an apparent density of 1.13 ± 0.02 mg/mm^3^, closely approximating the predicted density of pure LNG^21^ **(Fig. S1A)**. To determine whether the compacted LNG formulations exhibit sustained release profiles, we characterized their *in vitro* release kinetics under accelerated conditions–pH 7.4 phosphate-buffered saline supplemented with 3% w/w SDS surfactant, at 37°C and constant agitation (**Fig. 1E**, **1F**). Surfactant supplementation enhanced the solubility of LNG in the release medium, enhancing and accelerating dissolution due to LNG’s poor water solubility.

The *in vitro* drug release profiles revealed rapid dissolution of the 1×1 mm cylindrical formulations within 1 day, whereas the larger 2×2 mm and 4×4 mm formulations exhibited sustained release over 200 days **(Fig. 1E)**. The drug release rates for the 2×2 mm and 4×4 mm formulations were proportional to the formulation surface area, with the 4×4 mm formulations exhibiting a ∼4-fold greater initial surface area and a corresponding ∼4-fold higher daily release rate **(Fig. 1F)**. Notably, the release rates remained consistent throughout the experiment. These observations suggest that drug release from the compacted LNG formulations is governed by a surface area-controlled dissolution mechanism, and that increasing the formulations size can extend the release duration. Compacted monolithic LNG formulations manufactured using different tablet compression pressures, ranging from 159.2 MPa to 1114.1 MPa, had comparable bulk densities **(Fig. S1A)** and *in vitro* drug release rates **(Fig. S1B)**, indicating that the formulations were fully compacted across all pressures tested.

We further investigated whether compacted formulations of other poorly water-soluble drugs exhibit similar release kinetics. Using the same approach, we fabricated compacted formulations of quinestrol, an estrogen used in menopausal hormone therapy, and ivermectin, an antiparasitic for onchocerciasis and scabies. These formulations demonstrated excellent chemical stability when incubated at simulated physiological conditions (pH 7.4 PBS at 37°C and 60°C supplemented with surfactant) for up to 2 weeks, and exhibited sustained drug release under accelerated physiological conditions **(Fig. S2, S3)**. These results suggest that compacted monolithic formulations of various low-solubility small molecules drugs also undergo surface-mediated dissolution under physiological conditions, resulting in sustained drug release.

### Compacted levonorgestrel formulations provide sustained drug release for over 1 year following subcutaneous implantation in rats

To evaluate *in vivo* drug release kinetics of the compacted LNG formulations, cylindrical formulations of three different sizes (1×1 mm, 2×2 mm, and 4×4 mm, corresponding to surface areas of 4.71 mm^2^, 18.85 mm^2^, and 75.4 mm^2^, respectively) were individually implanted into the subcutaneous tissue of healthy Sprague Dawley rats **(Table S1)**. Plasma drug concentrations were measured weekly over a period exceeding one year **(Fig. 2A)**. Pharmacokinetic profiles for animals implanted with the largest (4×4 mm) formulation demonstrated consistent plasma drug levels with no discernible C_max_ or T_max_. Plasma drug concentrations were maintained above the equivalent human therapeutic drug level of 1 ng/mL in rats throughout the 53-week study **(Fig. 2B)**^22^. In contrast, plasma LNG concentrations in animals implanted with the 1×1 mm or 2×2 mm formulations remained above the limit of quantification (LLOQ of 25 pg/mL) for up to 3 months. Normalization of plasma drug concentrations to the initial surface area of each formulation revealed a clear correlation between formulation surface area and systemic drug levels **(Fig. 2C)**. Together, these results indicated that drug release from the monolithic formulations is primarily governed by the bulk surface area, consistent with *in vitro* observations, and suggest that a formulation with a surface area greater than 75.4 mm^2^ is sufficient to achieve and maintain human therapeutic drug levels in Sprague Dawley rats (∼1 ng/mL)^22^.

**Figure 2.**
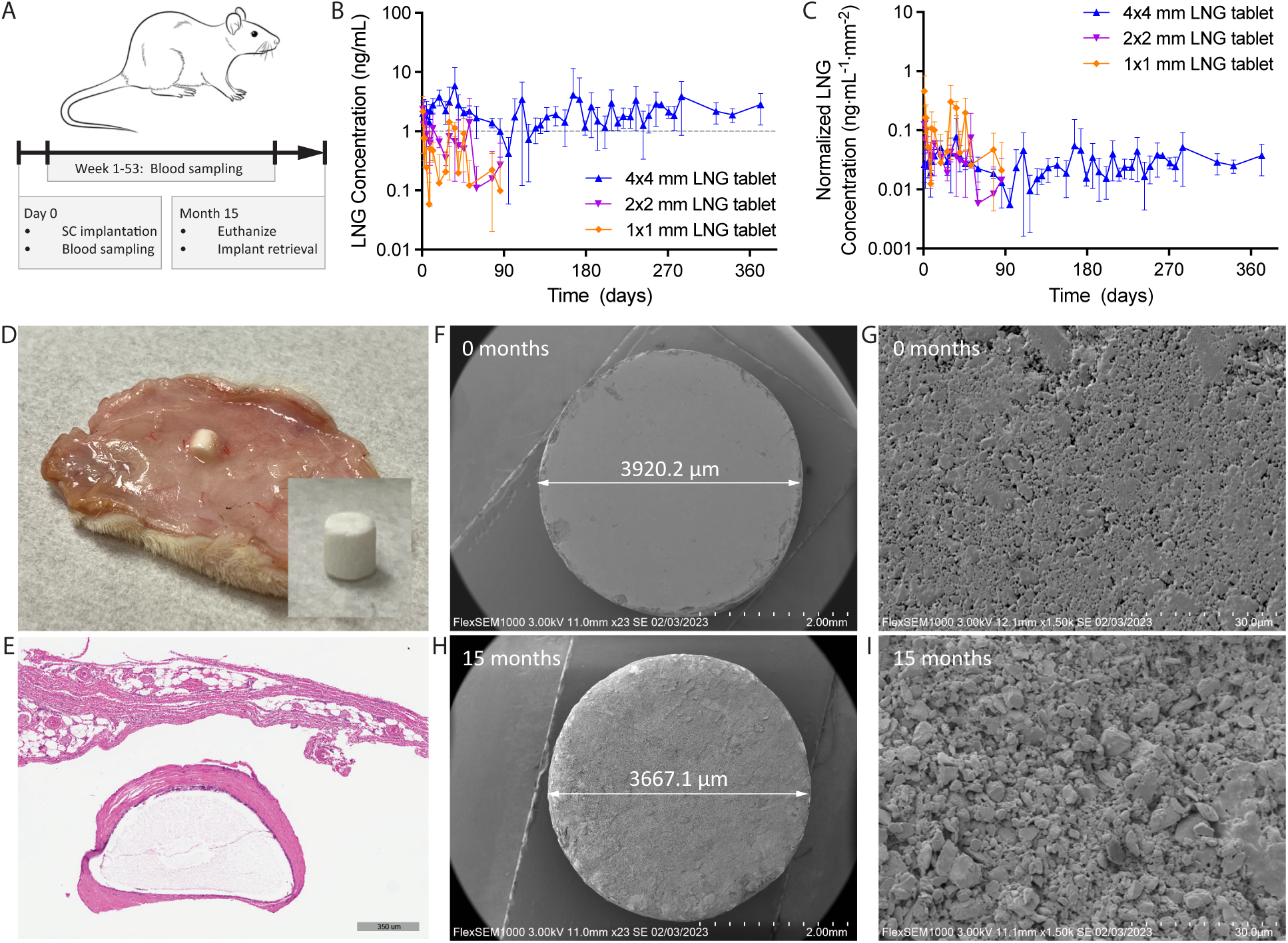
*In vivo* characterization of compacted LNG formulations. A) Experimental timeline. The formulations were implanted into the subcutaneous tissue on the dorsal region of Sprague Dawley rats. Blood was sampled over the course of a 53-week period. Animals were euthanized and the compacted LNG formulations were retrieved 15 months post implantation. B) Plasma concentration-time curves showing plasma LNG levels of 3 different formulation sizes over a 53-week period. Data plotted as mean ± SD (n=4). C) Normalized plasma concentration-time curves showing plasma LNG levels of 3 different formulation sizes that are normalized to the initial surface area of the formulations over a 53-week period. Data plotted as mean ± SD (n=4). D) Photograph of explanted 4×4 mm compacted LNG formulation. Inset: formulation was extracted from the subcutaneous tissue. E) H&E-stained tissue section showing the cross section of the compacted LNG formulation within the subcutaneous tissue. F) SEM image showing the top-down view of a 4×4 mm compacted LNG formulation prior to implantation. G) SEM image showing the surface of a 4×4 mm compacted LNG formulation prior to implantation. H) SEM image showing the top-down view of a 4×4 mm compacted LNG formulation 15 months post implantation. I) SEM image showing the surface of a 4×4 mm compacted LNG formulation 15 months post implantation.

The 4×4 mm compacted LNG formulations remained intact, solid, and structurally stable within the subcutaneous tissue after 15 months. The formulations retained sufficient mechanical integrity to allow for physical extraction and manipulation (**Fig. 2D**, **Fig. S4, Supplementary Video 1**). Histological analysis of the implantation site showed no evidence of active inflammation and revealed the formation of a defined fibrous capsule, ranging from 20.8 µm to 130.84 µm in thickness, surrounding the formulation–indicating good biocompatibility (**Fig. 2E**).

Detailed SEM analysis comparing the monolithic LNG formulation before implantation **(Fig. 2F,G)** and after 15 months *in vivo* **(Fig. 2H,I)** revealed a reduction in diameter from 3920.3 ± 0.3 µm to 3664.8 ± 35.6 µm, and a mass loss of 5.14 ± 0.83 mg, corresponding to an estimated surface erosion rate of 102.2 ± 14.3 µm per year. SEM imaging of the formulation surface prior to implantation **(Fig 2G)** and following retrieval **(Fig. 2I)** demonstrated that the tablet remained densely compacted with no significant development of enlarged internal porosity, supporting the hypothesis that drug partitioning occurs predominantly at the external surface rather than the internal porosity of the compacted meso-crystalline structure.

### Design of monolithic absorbable implant architectures

A common limitation in dissolution-controlled solid formulations–including aqueous crystalline suspensions and the compacted monolithic systems described above–is the gradual tapering of drug release rates over time, as the formulation size and surface area decreases during dissolution **(Fig. 1B)**. To achieve sustained therapeutic levels over a target duration, this tapering necessitates initially elevated drug release rates, resulting in supratherapeutic plasma drug levels early in the treatment. This phenomenon also results in a prolonged subtherapeutic phase after the treatment period. For instance, users of Sayana Press® injectable contraceptive may experience delayed return to fertility lasting over a year, likely due to unpredictable pharmacologic effects of prolonged subtherapeutic drug exposure^23^. Analogously, prolonged subtherapeutic exposure to antibiotics, anti-parasitics, anti-retrovirals, and chemotherapeutics may promote drug resistance and treatment failure^24–28^. Moreover, this release profile leads to inefficient drug utilization, reducing the achievable therapeutic duration of space-constrained dosage forms. An optimal long-acting drug delivery system would therefore provide consistent zero-order release, maintaining plasma drug levels at or slightly above the therapeutic threshold throughout the implant’s lifetime, followed by a rapid and complete cessation of drug release.

Drug release modulation from solid dosage forms have been extensively studied^29–31^, particularly in oral extended-release polymer matrix tablets^32–37^. Several systems capable of zero-order drug release over several hours in the gastrointestinal tract have been developed by combining insights in drug diffusion, polymer matrix erosion, and clever geometric designs^31,32,35,37^. Building on these principles, we adapted this strategy for overcoming inefficient drug utilization and dose tapering of compacted monolithic formulations for long-acting parenteral delivery by embedding them into space-efficient biodegradable devices. These devices modulate the exposed formulation surface area over multi-year timeframes, enabling precise tuning of drug release kinetics, including zero-order drug release and rapid end-of-life termination (**Fig. 1B**). Additionally, orthogonal modulation of formulation thickness provides a mechanism to adjust the total duration of drug release. Unlike prior systems reported in the literature, our device designs aim to be fully bioresorbable and highly space-efficient to maximize therapeutic lifetime while enabling minimally invasive insertion using a hypodermic needle or trocar, and providing orthogonal control over drug release rates and duration. To our knowledge, this is the first demonstration of this approach in the parenteral environment, and using polymer-free extended-release drug formulations over multi-month to multi-year timescales.

Accordingly, we designed biodegradable monolithic implants to address three major challenges hindering the clinical translation of compacted LNG formulations: (1) Tapering drug release due to shrinking surface area, leading to prolonged subtherapeutic periods and requiring initial supratherapeutic exposure; (2) poor mechanical integrity of the compacted formulations, which are brittle and prone to fracturing during handling and residence in the body; and (3) the need for integration of multiple tablets into a single, mechanically robust, and compact device for minimally invasive insertion, if needed, retrieval should users choose to discontinue therapy.

The rod-shaped monolithic implant consists of a biodegradable casing encasing the perimeter of six discrete compacted LNG formulations, each rectangular in shape and measuring 1.8 mm x 4 mm x 2 mm (**Fig. 3A**, **3B, Table S1**). The solid LNG formulations remained stable throughout the vacuum compression molding process used to manufacture the implants (**Fig. S5**). Use of multiple discrete tablets, rather than a single high-aspect-ratio tablet, reduces the risk of mechanical fracture due to bending. Drug release from the device occurs through the exposed top and bottom surfaces of each compacted tablet, with the surrounding casing providing structural support.

**Figure 3.**
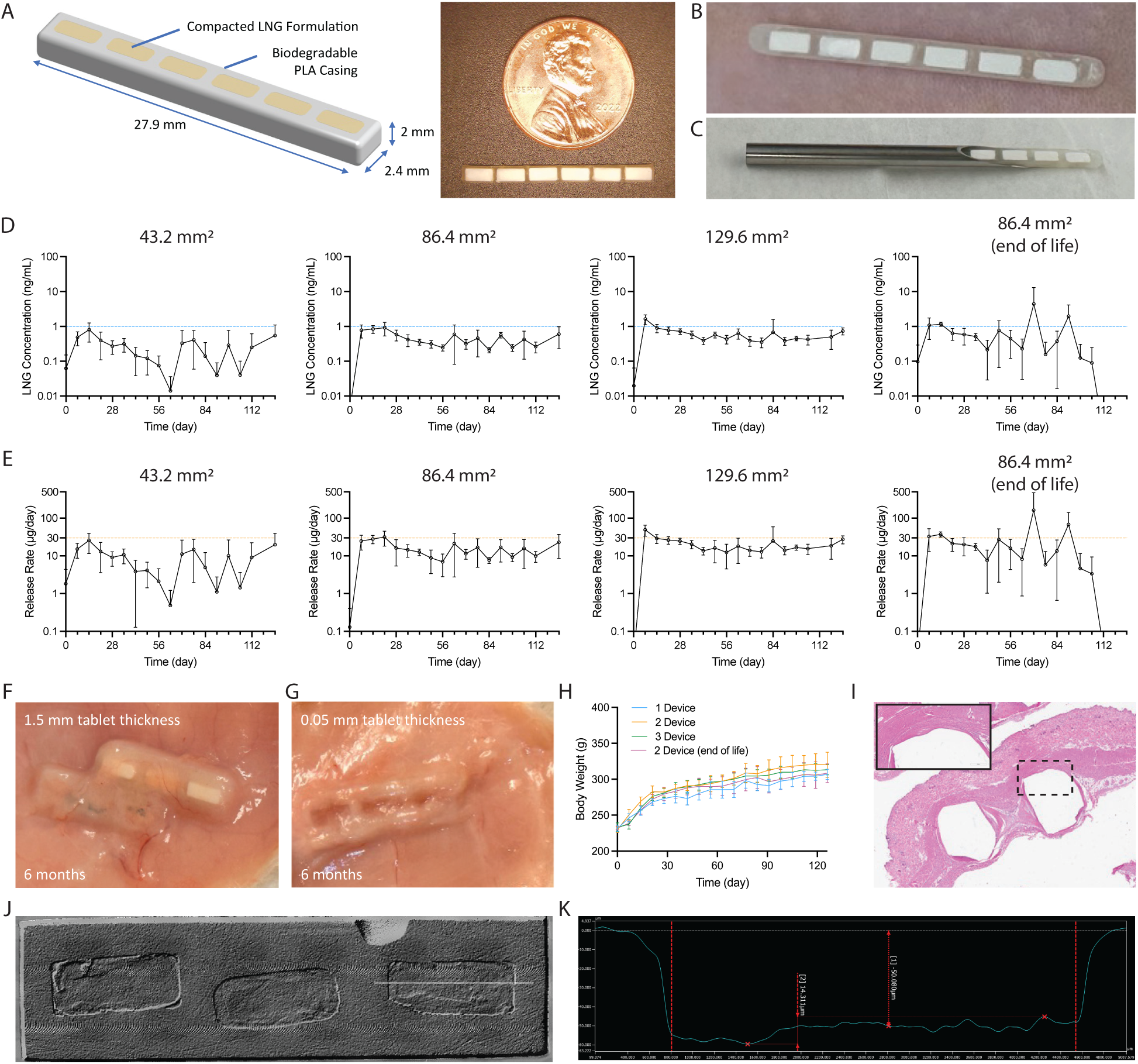
Monolithic LNG implants. A) Schematic and photograph of LNG-loaded MoSAIC implants. B) Photograph showing loading of a MoSAIC implant in a trocar for subdermal insertion. C) Photograph showing the ability to insert MoSAIC implants through an 8G trocar. D) Plasma LNG concentration-time curves of four different MoSAIC device designs over a 120-day period following subcutaneous implantation into the dorsal region of Sprague Dawley rats. Full-thickness devices were evaluated with 5 replicates (n=5). End-of-life devices were evaluated with 4 replicates (n=4). Data plotted as mean ± SD. E) Daily drug release rates for four different MoSAIC device designs over a 120-day period. Data plotted as mean ± SD F) Photograph a MoSAIC device loaded with 2 mm thick compacted LNG formulations 6-months after implantation in the subcutaneous tissue of a rat. G) Photograph a MoSAIC device loaded with 0.05 mm thick compacted LNG formulations 6-months after implantation in the subcutaneous tissue of a rat. H) Body weights of the rats receiving an implant over the course of the 120-day implantation period. Data plotted as mean ± SD (n ≥ 4). I) H&E-stained tissue section showing the cross section of the LNG-loaded MoSAIC devices within the subcutaneous tissue. J) Optical profilometry image of the top surface of a retrieved MoSAIC device, which was initially loaded with 2 mm thick compacted LNG formulations, 6 months after implantation in the subcutaneous tissue of a rat. The white line denotes the location of the measured cross-sectional profile. K) The measured profile along the length of the compacted LNG formulation. The distance along the length of device and the vertical height of the device, measured with respect to the top surface of the device, is plotted on the x-axis and y-axis, respectively.

These trocar-compatible device designs achieve an overall drug loading efficiency ≥68% by volume (**Table S1**), surpassing those of Jadelle® (∼35.5% by weight) and Nexplanon® (∼54.1% by weight). The overall implant diameter of 2.4 mm supports minimally invasive insertion via trocar, in line with Jadelle®, to ensure patient acceptability (**Fig. 3C**). The device architecture ensures a constant exposed formulation surface area throughout the therapeutic lifetime of the system, facilitating consistent drug release. Construction using high molecular weight poly-L-lactiv acid (PLLA), an FDA-approved biodegradable polyester, facilitates retrieval during use^38,39^ and complete degradation, erosion, and clearance from the body following depletion of encapsulated drug, eliminating the need for end-of-life surgical removal^40–43^.

To demonstrate mechanical durability under simulated use conditions, implants were inserted subcutaneously into *ex vivo* porcine tissue at a depth of 0.5 mm to mimic the clinical placement of Nexplanon®. An axial force of 60 N was applied to the skin surface directly above the implant to simulate maximum finger force encountered during routine use (**Fig. S6A**). Post-loading inspection revealed no evidence of implant damage, tablet fracture, or tablet displacement, confirming the implant’s structural integrity (**Fig. S6B**).

### Monolithic absorbable implants maintain consistent plasma drug levels following subcutaneous implantation in rats

To characterize *in vivo* drug release kinetics from monolithic LNG implants, 20 healthy Sprague Dawley rats were randomized into 4 experimental groups. Animals in the first three groups received 1, 2, or 3 devices, each containing 3 compacted monolithic tablets, corresponding to total exposed formulation surface areas of 43.2 mm^2^, 86.4 mm^2^, or 129.6 mm^2^, respectively. These ‘full-duration’ devices contained tablets that were 2 mm thick and are estimated to provide sustained drug release for ∼10 years based on previously determined erosion rates (**Fig. 1F**, **1H**). To evaluate end-of-life drug release kinetics, animals in the fourth group received 2 “end-of-life duration” devices loaded with 50 µm thick tablets, designed to release drug for ∼3 months. Plasma drug levels (**Fig. 3D**) and drug release rates (**Fig. 3E**) were measured weekly over a 4-month period.

In rats implanted with full-duration devices, plasma LNG levels remained stable throughout the study, consistent with zero-order release kinetics. Interestingly, despite previous results suggesting that an exposed formulation surface area of 75.4 mm^2^ was sufficient to maintain plasma LNG levels above the therapeutic threshold of 1 ng/mL, devices with 86.4 mm^2^ and 129.6 mm^2^ surface area exhibited subtherapeutic plasma concentrations (**Fig. 3D**) and drug release rates at approximately half of the 30 µg/day target (**Fig. 3E**). This discrepancy may stem from the presence of corners on the cylindrical compacted monolithic formulations, which may dissolve at a faster rate compared to flat surfaces, resulting in higher drug release rates. In the “end-of-life duration” implant group, plasma LNG levels remained similar to the “full-duration” for ∼3 months, followed by a rapid decline to undetectable levels (**Fig. 3D, E**). These results show a rapid cessation of drug release at the expected 3-month timepoint, supports the hypothesis that the system can deliver rapid return to fertility without a protracted sub-therapeutic phase.

To assess local tissue response, implants were retrieved 6 months post implantation. Gross inspection of the implants showed that full-duration devices remained intact with a substantial amount of drug remaining (**Fig. 3F**), while end-of-life devices were complete depletion of drug, leaving empty polymer casings (**Fig. 3G**), consistent with *in vivo* drug release data. Animals in all groups gained weight at a comparable rate over the study period, indicating good systemic tolerability **(Fig. 3H)**. Histological analysis of the implantation site revealed formation of a well-defined 30-260 µm-thick fibrous capsule and a resolved inflammatory response, indicating good biocompatibility (**Fig 3I**).

To verify the compacted LNG formulation erosion rate within the monolithic implants, devices were retrieved after 6 months, and their geometry was characterized using laser profilometry (**Fig. 3J**). A cross-sectional profile of the tablets (**Fig. 3K**) revealed uniform surface erosion with an average depth of ∼50 µm, closely matching previous estimates. Surface roughness analysis revealed a peak-to-trough variation of 15 µm, further supporting a uniform and consistent surface erosion mechanism.

LNG stability within the implants was also confirmed. Compacted LNG formulations from devices explanted after ∼1 year were extracted and analyzed for chemical purity. A ∼4% reduction in chemical purity was observed relative to freshly drug, indicating excellent long-term *in vivo* stability (**Fig. S7**).

Collectively, these results demonstrate that two full-duration, human-sized devices, with a combined exposed formulation surface area of 172.8 mm^2^, can be co-implanted in a manner analogous to the well-tolerated Jadelle® system to provide approximately 10 years of contraceptive protection. This system offers 2–3.3 fold longer coverage compared to Jadelle® and Nexplanon®, with the added advantage of smaller device size, full bioresorbability (eliminating the need for surgical retrieval), and rapid end-of-life drug release termination to support improved patient acceptability, adoption, and access.

Importantly, the monolithic implant architecture allows precise orthogonal control over drug release rate and duration by independently tuning formulation surface area and thickness. This capability enables a customizable family of implant designs tailored to a range of contraceptive durations aligned with individual user needs. For example, monolithic implants fitted with 0.1 mm, 0.2 mm, 1 mm, or 2 mm-thick compacted LNG formulations could provide coverage for 1, 2, 5, or 10 years, respectively.

### Design of monolithic shape-changing absorbable implant architectures which enable space-efficient tuning of drug release rates and therapeutic duration

The need for two separate implants to achieve therapeutic efficacy may limit patient acceptability and adoption. To address this, we developed a space-efficient, *in-situ* shape-changing monolithic implant architecture capable of providing therapeutic LNG release rates (i.e., 30 µg/day^22^, corresponding to an exposed formulation surface area of 172.8 mm^2^) from a single device (**Fig. 4A**). These designs increase the exposed formulation surface area multiple folds by dividing the original rod-shaped architecture into multiple planar sections or layers, at the cost of reducing formulation thickness, and thus, reduced therapeutic duration. Critically, this strategy preserves the constant exposed formulation surface area throughout the lifetime of the device, supporting consistent drug release rates and plasma drug levels.

**Figure 4.**
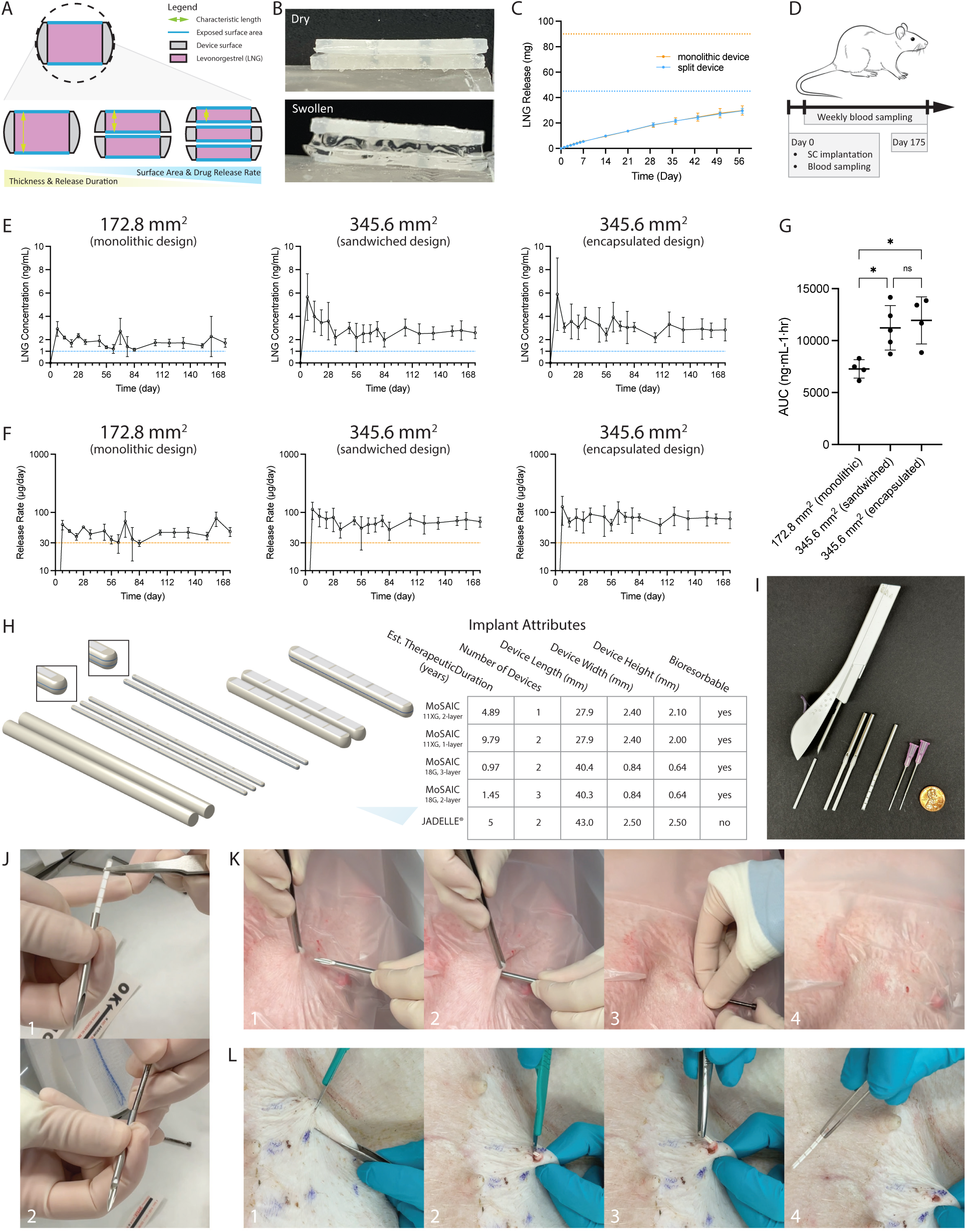
Space-efficient shape-changing monolithic LNG implant designs. A) Schematic showing the cross section of a MoSAIC implant design with one, two, or three layers. The compacted LNG formulation and PLLA casing are shown in pink and grey respectively. The exposed formulation surface area is denoted in blue, and the characteristic formulation thickness is denoted in green. Two- and three-layer designs are separated with a crosslinked xerogel that expands upon exposure to interstitial fluid to physically separate the layers. B) Photograph showing a two-layer shape changing MoSAIC design prior to (dry) and after (swollen) incubation in pH 7.4 PBS. C) *In vitro* drug release profiles for two MoSAIC device designs with equivalent exposed formulation surface area (i.e., two discrete a single layer MoSAIC devices and a single 2-layer shape-changing device). Devices were incubated in pH 7.4 PBS supplemented with 3% SDS at 37°C under sink conditions and constant agitation. Data plotted as mean ± SD (n = 5). D) Experimental timeline for *in vivo* pharmacokinetic evaluation of three different LNG-loaded MoSAIC device designs. Devices were individually implanted into the subcutaneous tissue on the dorsal region of Sprague Dawley rats. Each device design was evaluated with n = 5 replicates. Blood was sampled over a 175-day period. E) Plasma LNG concentration-time curves for 3 different MoSAIC device designs over a 175-day. Data obtained from n≥4 replicates and plotted as Mean ± SD. F) Daily drug release rates for 3 different MoSAIC device designs over a 175-day period. Data obtained from n≥4 replicates and plotted as Mean ± SD. G) AUC for the 3 different MoSAIC device designs following subcutaneous implantation in rats. Data obtained from n≥4 replicates and plotted as Mean ± SD. H) Models of four different MoSAIC implant designs and the commercial Jadelle® system for long-term reversible contraception. Estimated therapeutic durations are calculated from the measured *in vivo* dissolution rates of compacted LNG formulations in rats. I) Photograph showing the relative size of 2 different MoSAIC device designs, the commercial Nexplanon® system, and a model of the commercial Jadelle® system for long-term reversible contraception. J) Photographs showing loading of a 2-layer MoSAIC contraceptive device into a trocar for subsequent subcutaneous placement. K) Photographs showing the procedure for minimally invasive subcutaneous placement of a 2-layer MoSAIC contraceptive implant into the rear flank of an anesthetized pig. L) Photographs showing the procedure for minimally invasive retrieval of a 2-layer MoSAIC contraceptive implant from the rear flank of an anesthetized pig 2-weeks post insertion.

To overcome impeded drug release from opposed formulation surfaces, a dehydrated hyaluronic acid hydrogel layer was incorporated between the layers. Upon subcutaneous implantation, interstitial fluid induces hydrogel swelling, physically separating the layers and enabling unimpeded drug release (**Fig. 4B**). This design is highly space-efficient since the hydrogel layer increases the implant’s dry-state volume by <5% while preserving its narrow geometry, allowing for minimally invasive trocar-based insertion.

To evaluate whether the hydrogel layer affects drug release from the compacted formulations, we compared *in vitro* release kinetics from two systems with equivalent exposed formulation surface areas (**Fig. 4C, Table S1**). The first, a shape-changing MoSAIC device, consists of two 1 mm-thick planar sections separated by a swollen hydrogel layer, such that half the formulation surfaces oppose the hydrogel. The second control group includes two discrete, free-floating 2 mm-thick monolithic devices with all surfaces exposed to the release medium. Both systems exhibited identical drug release profiles, indicating that neither erosion nor drug release from the compacted LNG formulation surfaces was impeded by the presence of the hydrogel **(Fig. 4C)**.

To compare *in vivo* performance, healthy Sprague Dawley rats were randomized into three experimental groups. The first group received four 1-layer “half-length” monolithic devices with a total exposed formulation surface area of 172.8 mm^2^ (design 13, **Table S1**). The second group received four 2-layer “half-length” devices with a total exposed formulation surface area of 345.6 mm^2^ (design 15, **Table S1**) and where the two layers were either separated by a 0.1 mm hydrogel. The third group also received 2-layer “half-length” devices with a total exposed formulation surface area of 345.6 mm^2^ (design 13, **Table S1**), however these layers were fully encapsulated with a 0.1 mm hydrogel layer on all sides (design 14, **Table S1**). Plasma LNG levels were measured weekly for 6 months (**Fig. 4D**). All groups maintained plasma LNG concentrations above 1 ng/mL and drug release rates in excess of 30 µg/day throughout the study (**Fig. 4E**, **4F),** confirming that the extrapolated compacted formulation surface area of 172.8 mm^2^ is sufficient to achieve human-equivalent therapeutic levels in rats (**Fig. 3G**, **3H**). Notably, shape changing MoSAIC groups showed ∼2-fold higher plasma LNG levels, daily release rates, and AUC_0-4wks_ compared to the standard monolithic group, consistent with their doubled exposed formulation surface area (**Fig. 4G**). These findings support a surface-erosion based release mechanism and indicate that the hydrogel layer does not significantly impeded drug release.

Together, these results demonstrate a space-efficient, orthogonal method to independently modulate LNG release rates and therapeutic duration. Importantly, the shape-changing, multilayer MoSAIC design enables therapeutic LNG delivery from a single rod-shaped 2-layer implant for a projected ∼5 years (**Fig. 4H**, **4I, Table S1, Table S2**). The same principle can be extended to smaller-diameter designs suitable for insertion via standard 18-gauge hypodermic needles, enabling 1–1.5 years of contraceptive (**Fig. 4H, 4I, Fig. S8, Table S1**). Recognizing that the physicochemical properties, target therapeutic drug levels, and desired therapeutic durations of other long-acting drug therapies (e.g., ivermectin, quinestrol) differ from LNG, this generalized shape-changing approach offers a versatile platform for developing minimally invasive, biodegradable long-acting drug delivery implants for a wide range of chronic indications.

We further demonstrate the feasibility of minimally invasive insertion and retrieval of the full-length (human-sized) 2-layer MoSAIC contraceptive implant in a large animal porcine model, which better approximates human skin anatomy compared to rodents. The implant was loaded into a trocar and temporarily secured in place using a plunger (**Fig. 4J, Supplementary Video 2**). Placement into the subcutaneous tissue of the rear flank proceeded via trocar insertion and advancement (along with the implant), followed by trocar retraction while stabilizing the implant within the tissue with the plunger (**Fig. 4K, Supplementary Video 3**). The plunger was then withdrawn from the tissue, and the incision closed using bandages (i.e., Tegaderm®) or surgical adhesive. We further demonstrate the feasibility of removing the implant (**Fig. 4L, Supplementary Video 4**). At two weeks post-implantation, the device was retrieved by identifying and securing the implant through the skin, making a small incision at one end, and withdrawing the implant as a single intact, rigid object using forceps. The incision was then closed using bandaging or surgical glue.

## Discussion

Contraceptive drugs empower women to prevent unintended pregnancies and to space planned ones. Their effectiveness, however, hinges on the maintenance of consistent drug levels over time, making adherence to dosing regimens essential. To reduce the burden of frequent dose administration, long-acting implantable contraceptives have been developed and approved. These systems are among the most effective reversible contraceptive options available under real-use conditions, yet their requirement for surgical of implantation and removal limits accessibility, particularly in low-resource settings. Each year, an estimated 257 million women who wish to avoid pregnancy are not using modern contraceptive methods, underscoring the need for next-generation systems that are more accessible and acceptable^17^.

In response, we developed the MoSAIC system–a monolithic, shape-changing contraceptive implant that is smaller, less invasive, and/or longer lasting than current available systems such as Jadelle®. MoSAIC was specifically designed to overcome key barriers to user acceptability and treatment access. To improve user acceptability and discretion, we developed and employed space-efficient compacted LNG formulations and shape-changing device architectures to minimize overall implant size, thereby reducing subdermal palpability and visibility. This approach also minimizes invasiveness by enabling device insertion using standard 18G hypodermic needle for shorter-duration coverage (1–1.5 years), and reducing the number trocar-inserted devices needed for longer-term use. For example, MoSAIC enables 5-year contraception with a single device, compared to the two-device Jadelle® system. Critically, the system is constructed entirely from bioresorbable, FDA-approved materials, which eliminates the need for end-of-life surgical removal–improving treatment access, particularly in low-resource settings where trained medical personnel and clinical facilities are limited.

MoSAIC’s unique drug release kinetics minimizes risk of side effects and enables longer-term coverage compared to existing systems. The system’s zero-order drug release profile maximizes drug utilization, enabling up to 10 years of coverage and significantly reducing the lifetime number of devices and procedures a user might require compared to Jadelle®. It also minimizes plasma concentration fluctuations (i.e., large peak to trough plasma drug levels), although LNG’s broad therapeutic window makes it less susceptible to such variation. The absence of dose tapering and sharp end-of-life cessation of drug delivery mitigates the risk of long-term subtherapeutic drug exposure that often leads to delayed return to fertility–a concern with extended release formulations like Depot Provera^®^ and Sayana Press^®^.

These capabilities were enabled by two key advances: the development of compacted, extended-release LNG formulations with exceptionally high drug loading efficiency, and the design of a flexible, space-efficient, structurally robust multilayer shape-changing device architecture that enables independent control of both drug release rate and duration. We demonstrated that compacted monolithic formulations of LNG and other low-solubility drugs exhibit gradual surface-mediated dissolution in aqueous environments, even in the presence of internal porosity. This behavior, reminiscent of the micronized drug particles used in long-acting injectable crystalline suspensions, likely arises from limited water penetration into the formulation core due to the hydrophobic nature of LNG (i.e., low surface energy limits wetting), poorly connected internal pores, and/or slow transport of solubilized drug from within the tortuous internal pore network leading to saturation-limited partitioning of LNG in the core. While further mechanistic studies are needed, this behavior enables the scalable, low-cost manufacturing of millimeter-scale formulations with both extremely high drug loading (∼100% w/w) and multi-year release durations–well beyond the capacity of injectable aqueous suspensions and conventional polymer-matrix systems.

By incorporating these formulations into the MoSAIC platform, we created a system in which drug release is confined to two planar surfaces. This architecture allows for independent tuning of drug release rate and duration by adjusting surface area and formulation thickness, respectively. Because the exposed surface area remains constant, dissolution proceeds with predictable zero-order kinetics. The layered, shape-changing design further allows for flexible control of these parameters within a prescribed implant footprint. Using this strategy, we demonstrate three versions of the MoSAIC implant capable of providing therapeutic LNG delivery for different durations: one capable of providing up to 10 years of contraception using 2 devices; another capable of delivering up to 5 years of coverage with a single device; and a third, ultra-thin two-device system compatible with 18G needle insertion and capable of sustaining drug release for 1 year.

Overall, the MoSAIC contraceptive system represents a significant advancement over existing long-acting reversible contraceptive implants. It is smaller, less invasive, and/or longer lasting compared to existing implants and offer users enhanced access and user experience, particularly in low resource settings. More broadly, the exceptional drug loading efficiency, ultra-long drug release profiles, and fully bio-resorbable nature of the MoSAIC platform has the potential to shift the current standard of chronic drug therapy from a paradigm of frequent and repeated dose administration, which is associated with suboptimal adherence, to a “dose it and forget it” approach where dose adherence is engineered into the system.

While this study focused on the delivery of LNG for long-acting contraception, the same system (and drug formulation) may support non-contraceptive indications such as dysmenorrhea, menorrhagia, endometriosis, and polycystic ovary syndrome (PCOS). The MoSAIC platform is also generalizable to other long-acting therapeutic applications. The modular nature of the device allows for tuning of layer count, formulation thickness, and formulation surface area to accommodate the dissolution and drug release kinetics of compacted formulations of other potent, low-solubility drugs. Potential indications include menopausal hormone therapy, breast or prostate cancer (e.g., quinestrol and estradiol), mass drug administration campaigns for parasitic infections such as onchocerciasis or malaria (e.g., ivermectin), HIV pre-exposure prophylaxis (PrEP), and chronic psychiatric conditions such as schizophrenia and bipolar disorder (e.g., pimozide and aripiprazole). Interestingly, the modular nature of this platform further enables easy delivery of drug combinations (e.g., contraception and HIV PrEP). In each case, treatment-specific drug release rates and durations could be achieved through appropriate structural modifications.

Despite its promise, the MoSAIC platform has important limitations. It is best suited for the delivery low-solubility, high-potency, and chemically stable active pharmaceutical ingredients. Drug selection must carefully consider both the limited volume available in the minimally invasive implants and the intended therapeutic duration. Candidate compounds must also remain stable under physiological conditions (37°C, interstitial fluid) for the full duration of delivery. Moreover, while the MoSAIC architecture affords flexibility in geometry and release kinetics, successful implementation requires the drug be compacted into solid monolithic formulations and capable of achieving therapeutic partitioning rates from the exposed surface area. These constraints pose challenges for the delivery of biologics, water-soluble drugs, or drugs requiring large doses. Finally, although we demonstrated minimally invasive insertion, retrieval, biocompatibility, and sustained delivery of LNG at human-equivalent levels in animal models, additional preclinical and clinical studies will be essential to support translation into human use.

## Materials and Methods

Dulbecco’s Phosphate-Buffered Saline (PBS) was purchased from Gibco by Life Technologies (Woburn, USA). Levonorgestrel (LNG, CAS# 797-63-7) was purchased from Austin Chemical, Inc. (sourced from Biosynth Carbosynth, COO, China). Ivermectin (CAS# 70288-86-7) was purchased from MedChemExpress. Quinestrol (CAS# 152-43-2) was purchased from Santa Cruz Biotechnology, USA. Sodium dodecyl sulfate (SDS, CAS# 151-21-3) was purchased from Thermo Fisher Scientific (28312). 2-hydroxypropyl beta cyclodextrin (HP-β-CD, CAS 128446-35-5) was purchased from Sigma, USA (778966). Poly (L-lactic acid) (PLLA, Mn 197,641, PDI 1.69) was purchased from Akina Inc., USA (AP008). Hyaluronic acid (300-500 kDa MW, 600-01-07) was purchased from Contipro a.s. (Czech Republic).

### Development of compacted drug formulations

Solid formulations composed of 100% w/w drug was manufactured by tablet compaction of as-received lyophilized product using an RD10A Natoli Tablet press (St. Charles, MO, USA). In a typical experiment compacted LNG formulations were formed using a compaction pressure of 208.3 MPa, although a broader range of pressures ranging between 39.7 MPa and 636.6 MPa were also explored. Magnesium stearate, a common lubricant used in tableting, was excluded from this process. The solid formulations were stored at 4°C and protected from moisture until use. Tablet density was calculated by dividing the gross weight of the formulation by its volume, as determined using optical profilometry.

### *In vitro* LNG release kinetics of compacted formulations

To characterize the drug release profile of compacted drug formulations, formulations were individually incubated in pH 7.4 PBS supplemented with 3% SDS or 10% HP-β-CD at 37°C, sink conditions, and under constant agitation^44^. At predetermined timepoints, 1.0 mL of release medium was sampled and replaced. Drug concentrations within the samples were analyzed using an Agilent 1260 Infinity I HPLC equipped with a UV detector, and an Agilent Poroshell 120 EC-C-18 2.7 µm 3.0 x 50 mm column (Agilent 699975-302). 5 µL of sample was loaded onto the column, heated at 50°C, using a mobile phase consisting of 5% acetonitrile and 95% of 0.1% formic acid. Gradient elution was carried out over a 4.5-minute period with a flow rate of 1.0 mL/min starting at 5% of acetonitrile and 95% of 0.1% formic acid and ending at 95% of acetonitrile and 5% of 0.1% formic acid. LNG was detected using a UV absorbance of 250 nm. LNG concentration was determined as the area under the main peak.

### Synthesis of methacrylated hyaluronic acid

2 g of hyaluronic acid was dissolved in 100 mL of water and stirred at 500 rpm for 2 hours at room temperature. 1.6 mL of methacrylic anhydride was added dropwise to the solution. 5 M NaOH was added dropwise to the solution to adjust the final pH to 8.5. The solution was protected from light and stirred at 4°C for 24 hours. 2.92 g of NaCl was added to the solution followed by precipitation of the polymer in ethanol. The resulting product was then isolated by centrifugation and washed in ethanol. The product was then redissolved in DI water and dialyzed against DI water for 3 days at 4°C.

### Manufacturing of monolithic and MoSAIC implants

Monolithic implants were fabricated using a vacuum compression molder. First, a positive implant casing mold was designed using SolidWorks and printed out of Rigid 10K resin and HTL resin using a Form 3 printer (Formlabs, USA) or BMF printer (Boston Micro Fabrication, USA), respectively. These were then used to cast a two-part separable negative PDMS mold for subsequent vacuum compression molding. PLLA was introduced into the PDMS molds and heated at 120°C for 10 minutes while under vacuum. The PLLA casing was cooled to room temperature and separated from the PDMS mold. Compacted drug formulations were manually inserted into the PLLA casing, heated at 120°C for 10 minutes while under vacuum, and cooled to room temperature. The monolithic devices were stored at 4°C and protected from moisture and light until use.

MoSAIC implants were constructed by stacking two or more monolithic implants alongside each other in a silicone mold and separated 0.1 mm apart. A stock photo-initiator solution was prepared by dissolving 60 mg of Irgacure I2959 in 3.6 mL of DI water and heated at 65°C for 12 hours while protected from light. A polymer stock solution was prepared by dissolving 200 mg of methacrylated hyaluronic acid and 4 mg of N,N’-methylenebisacrylamide in 1.6 mL of pH 7.4 PBS. The solution was heated at 37°C for 4 hours. To this stock, 2.4 mL of photo-initiator stock was added, vortexed, and incubated at 37°C for 15 minutes. 4 µL of 25% glutaraldehyde solution was added to the solution and vortexed. The homogenous solution was then centrifuged at 1000 rpm for 3 minutes to remove air bubbles. 100 µL of this polymer solution was dispensed into each mold, between the monolithic devices, and allowed to dry overnight at room temperature. The devices were subsequently exposed to UV irradiation in a UV oven for 15 minutes to facilitate UV-initiated hydrogel crosslinking. The MoSAIC devices were stored at 4°C and protected from moisture and light until use.

### *In vitro* characterization of LNG stability

The *in vitro* stability of levonorgestrel formulations was evaluated prior to and after integration into the MoSAIC implants using reverse phase HPLC. The samples were subjected to vacuum compression molding process whereby formulations are heated at 200°C for 30 minutes. LNG was physically extracted from the devices, weighed, and dissolved in 50% acetonitrile and 50% water at a concentration of 0.5 mg/mL. The chemical purity of LNG within the samples was measured using RP HPLC. Samples were analyzed using an Agilent 1260 Infinity I HPLC equipped with a UV detector, and an Agilent Poroshell 120 EC-C-18 2.7 µm 3.0 x 50 mm column (Agilent 699975-302). 5 µL of sample was loaded onto the column, heated at 50°C, using a mobile phase consisting of 5% acetonitrile and 95% of 0.1% formic acid. Gradient elution was carried out over a 4.5-minute period with a flow rate of 1.0 mL/min starting at 5% of acetonitrile and 95% of 0.1% formic acid and ending at 95% of acetonitrile and 5% of 0.1% formic acid. LNG was detected using a UV absorbance of 250 nm. LNG concentration was determined as the area under the main peak. LNG purity was determined as the ratio of the area under the main LNG peak and the total area under all peaks.

### Mechanical characterization of MoSAIC implants

The structural integrity of MoSAIC implants under simulated use conditions was evaluated in *ex vivo* porcine tissue. A freshly excised 1-inch-thick porcine tissue section was harvested from the flank and mounted to a rigid steel support. Full length devices were inserted into the tissue at a depth of 0.5 cm from the skin surface. A 60 N point load, which is the estimated maximum force that can be exerted by a finger, was applied overtop of the center of the device using a 1/8” diameter steel plunger fastened to the upper gripper of an Instron testing machine equipped with a 500 N load cell. Displacement was applied to the specimen at a rate of 15 mm/min until a force of 60 N was achieved. Following loading, the device was retrieved from the *ex vivo* tissue and examined for mechanical failure.

### *In vitro* characterization of LNG release kinetics from MoSAIC implants

To characterize the drug release profile of compacted LNG formulations, formulations were individually incubated in pH 7.4 PBS release medium supplemented with 10% HP-β-CD at 37°C, sink conditions, and under constant agitation^44^. At predetermined timepoints, 1.0 mL of release medium was sampled and replaced. Release medium was exchanged at regular intervals to maintain physiological pH. Drug concentrations within the samples were analyzed using an Agilent 1260 Infinity I HPLC equipped with a UV detector, and an Agilent Poroshell 120 EC-C-18 2.7 µm 3.0 x 50 mm column (Agilent 699975-302). 5 µL of sample was loaded onto the column, heated at 50°C, using a mobile phase consisting of 5% acetonitrile and 95% of 0.1% formic acid. Gradient elution was carried out over a 4.5-minute period with a flow rate of 1.0 mL/min starting at 5% of acetonitrile and 95% of 0.1% formic acid and ending at 95% of acetonitrile and 5% of 0.1% formic acid. LNG was detected using a UV absorbance of 250 nm. LNG concentration was determined as the area under the main peak.

### Pharmacokinetics of compacted LNG formulations in rats

All animal experiments were approved by and performed in accordance with the Committee on Animal Care at MIT. 200-225 g female Sprague Dawley (SAS SD strain 400) were purchased from Charles River. Compacted LNG formulations and LNG-loaded MoSAIC implants were implanted into the subcutaneous tissue of rats using a trocar. At predetermine timepoints, blood was sampled from the lateral tail vein and collected in EDTA microtainer capillary blood collection tubes (Becton Dickinson, BD365974-MI). Plasma samples were extracted via centrifugation at 2000 g and stored at -80°C. LNG concentrations within the plasma samples were measured by triple quadrupole liquid chromatography-tandem mass spectrometry (LC-MS/MS).

Non-compartmental pharmacokinetic analysis was used to calculate the daily drug release rates from each system (equation 1). Significant changes in the body weight of the rats were observed over the course of the experiment resulting in a change in drug clearance over time (**Fig. 3H**). The predicted LNG clearance in the rats was calculated by bodyweight-dependent allometric scaling^45^ based on clearance values reported by Ko et al.^22^ (i.e., CL_IV_ = 1.01 L/hr in 305 g Sprague Dawley rats).

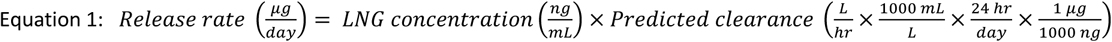

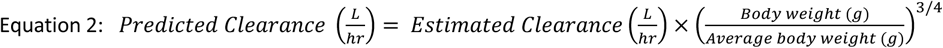

### Biocompatibility of compacted LNG formulations and MoSAIC devices in rats

Compacted LNG formulations and LNG-loaded MoSAIC devices were implanted into the subcutaneous tissue of 200-225 g female Sprague Dawley rats for up to 15 months. The implants and the surrounding tissue was subsequently excised, grossed, and fixed in 10% neutral buffered formalin for 24 hours, and stored in 70% ethanol. Tissue was embedded within paraffin blocks, sectioned into 5-micron tissue sections, and stained with hematoxylin and eosin. The inflammatory response and degree of fibrosis surrounding the implants was reviewed by a veterinary pathologist (Dr. Bronson, The Hope Babette Tang Histology Facility, Swanson Biotechnology Center, MIT).

### Minimally invasive implant insertion and retrieval in pigs

All animal experiments were approved by and performed in accordance with protocols approved by the Committee on Animal Care at the Massachusetts Institute of Technology. We demonstrate minimally invasive insertion and retrieval of the MoSAIC implants in a large animal model (75-kg Yorkshire swine; Cummings School of Veterinary Medicine at Tufts University, Grafton, MA). This model was used because its skin and subcutaneous tissue anatomy is similar to that of humans. Animals were fasted overnight before procedures to ensure safe anesthesia and to avoid aspiration. Pigs were sedated with Telazol (5 mg/kg; tiletamine/zolazepam) and xylazine (2 mg/kg) or dexmedetomidine (0.03 mg/kg) and midazolam (0.25 mg/kg), intubated, and maintained on 1 to 3% isoflurane in oxygen. Their heart rate, respiratory rate, end tidal CO_2_, SpO_2_, and temperature were monitored while anesthetized.

Sterile full-length 2-layer LNG-loaded MoSAIC devices were loaded into 10-gauge stainless steel trocars and inserted into the subcutaneous tissue at the rear flank of the pigs, along a superficial plane beneath the skin, on day 0 and day 7. A steel plunger was used to maintain the location of the implant within the subcutaneous tissue while the trocar was retracted. The plunger was then retracted, and the small incision was closed using surgical glue and bandaged using Tegaderm® transparent film dressing. On day 14, the implants were retrieved by forming a small surgical incision at one of the MoSAIC devices, grasping the exposed device end using forceps, and extraction of the entire device from the subcutaneous tissue.

### *Ex vivo* characterization of LNG formulations and MoSAIC implants

The weight and geometry of compacted LNG formulations and MoSAIC implants was measured using a Mettler Toledo XSR205 analytical balance and a Keyence VK-X3000 3D surface profiler, respectively. Formulation density was calculated as *ρ* = *mass* (*mg*)/*volume* (*mm*^3^). The *in vivo* erosion rate for the compacted LNG formulations were calculated from the change in formulation thickness over the total time of implantation (i.e., 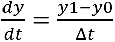 where y is the formulation thickness at a specified timepoint). The surface morphology of the compacted LNG formulations, before and after subcutaneous implantation in rats, was evaluated using a Hitachi FlexSEM TM-1000 II (SU1000, Tokyo, Japan) scanning electron microscope. The chemical stability of LNG within MoSAIC implants prior to and following implantation in the subcutaneous tissue of rats for a period of 1 year was evaluated using RP HPLC in an analogous manner described above.

### SEM Sample Preparation and Imaging Conditions

Prior to vacuum pump down, the LNG formulations and MoSAIC implants were coated with approximately 15 nm of gold using JEOL USA’s Smart Coater (Peabody, MA USA). This conductive coating prevented excessive surface charging artifacts in images. The samples were mounted using double sided carbon tape (Ted Pella Inc) and imaged in high vacuum mode using the secondary electron (SE) detector from Hitachi FlexSEM TM-1000 II (Tokyo, Japan). Low voltage imaging (3 kV) was used to prevent damage from electron bombardment, but 3 kV also provided high surface detail. Typical imaging conditions would also include a spot intensity of 50 (based unitless scale from 1 - 100) and a working distance between 6.5 - 8 mm.

### Statistical analysis

Unpaired two-sided Student’s t-tests and one-way analysis of variance (ANOVA) with Tukey’s multiple comparisons tests were performed using GraphPad Prism (Version 9.4.1). A value of P<0.05 was considered statistically significant. Figure captions and text describe the number of replicates used in each study. Figure captions define the center line, and error bars presented in the plots.

## Acknowledgments

The authors are grateful for constructive discussions with Dr. Robert Langer, Dr. Giancarlo Francese, Dr. Dennis Lee, Dr. Bret Berner, Dr. Jeremy Blum, Dr. Steve Zale, Dr. Victor Nguyen, and Dr. Mark Milad. The authors also thank Kathy Cormier at The Hope Babette Tang Histology facility in the MIT Koch Institute Swanson Biotechnology Center for technical support, Dr. Roderick Bronson for his expertise in providing histological assessment, and Virginia Fulford for providing the illustrations in Figure 1.

## Author contributions

J.L. and G.T. conceived of the concept and designed all experiments. J.L., B.G.C., P.K., C.C., J.Y.L., S.R.L., Y.S., K.J.Q., F.C., E.D., I.T.T., J.E.G., K.M., J.D.B., S.Y.O. developed, manufactured, and characterized all formulations. J.L., A.L., B.N.M., developed characterization methods and characterized all formulations. J.L., B.G.C., P.K., C.C., J.Y.L., S.R.L., Y.S., K.J.Q., F.C., E.D., I.T.T., J.E.G., K.M., J.D.B., S.Y.O. designed, manufactured, and characterized all devices. J.L., B.G.C., P.K., C.C., J.Y.L., S.R.L., Y.S., K.J.Q., F.C., E.D., I.T.T., J.E.G., A.M.H., N.F., A.P., K.S., B.L., M.P., A.G., performed *in vivo* characterization of formulations and devices. J.L., B.G.C., P.K. J.Y.L., E.D., A.L., B.M., J.D.B., S.Y.O., P.R.C., R.L. analyzed all data. J.L., G.T., R.L. prepared figures and wrote the manuscripts with edits from all authors. J.L. and G.T supervised all aspects of the work.

## Data and materials availability

All data are available in the main text or the supplementary materials. The data used to generate the figures and numerical statistics in this paper can be found as attached and in the Supplementary Information.

