## Supplemental Information for "Monolithic Shape-Shifting Absorbable Implants for Long-Term Contraception"

**Table S1:** Specifications of compacted formulations and MoSAIC devices

| | Formulation /Device | API[a] | Formulation Dimensions (mm)[b] | Device Dimensions (mm) | Drug Loading (mg) | Drug Loading Efficiency (% vol) | Exposed Formulation Surface Area ($mm^2$) | Estimated Therapeutic Duration (years) |
|---|---|---|---|---|---|---|---|---|
| **Compacted Formulations** | | | | | | | | |
| **1** | Cylindrical | LNG | 1.0 × 1.0 (D×H) | - | 0.83 ± 0.07 | 100 % | 4.71 | - |
| **2** | Cylindrical | LNG | 2.0 × 2.0 (D×H) | - | 7.45 ± 0.65 | 100 % | 18.85 | - |
| **3** | Cylindrical | LNG | 4.0 × 4.0 (D×H) | - | 61.78 ± 3.58 | 100 % | 75.40 | - |
| **4** | Rectangular | LNG | 4.0 × 1.8 × 2.0 (L×W×H) | - | 15.22 ± 1.30 | 100 % | 37.60 | - |
| **5** | Cylindrical | Quinestrol | 3.0 × 3.0 (D×H) | - | 26.73 ± 3.95 | 100 % | 42.41 | - |
| **6** | Cylindrical | Ivermectin | 3.0 × 3.0 (D×H) | - | 28.22 ± 1.75 | 100 % | 42.41 | - |
| **Full-length MoSAIC devices** | | | | | | | | |
| **7** | one 1-layer trocar device | LNG | 6 tablets 4.0 × 1.8 × 2.0 (L×W×H) | 27.9 × 2.4 × 2.0 (L×W×H) | 91.32 ± 7.80 | 73.3% | 86.4 | NA |
| **8** | one 2-layer trocar device | LNG | 12 tablets 4.0 × 1.8 × 1.0 (L×W×H) | 27.9 × 2.4 × 2.1 (L×W×H) | 91.32 ± 7.80 | 68.0% | 172.8 | 4.89 |
| **9** | two 1-layer trocar devices | LNG | 12 tablets 4.0 × 1.8 × 2.0 (L×W×H) | 27.9 × 2.4 × 2.0 (L×W×H) | 91.32 ± 7.80 | 73.3% | 172.8 | 9.78 |
| **10** | two 2-layer trocar devices | LNG | 24 tablets 4.0 × 1.8 × 1.0 (L×W×H) | 27.9 × 2.4 × 2.1 (L×W×H) | 91.32 ± 7.80 | 68.0% | 345.6 | 4.89 |
| **11** | two 3-layer 18G devices | LNG | 36 tablets 6.20 × 0.39 × 0.20 (L×W×H) | 40.4 × 0.84 × 0.64 (L×W×H) | 19.62[c] | 61.6% | 175.3 | 0.97 |
| **12** | three 2-layer 18G devices | LNG | 36 tablets 6.20 × 0.39 × 0.30 (L×W×H) | 40.4 × 0.84 × 0.64 (L×W×H) | 29.43[c] | 61.6% | 175.3 | 1.45 |
| **Half-length MoSAIC devices (in vivo rats)** | | | | | | | | |
| **13** | one 1-layer trocar device | LNG | 3 tablets 4.0 × 1.8 × 2.0 (L×W×H) | 13.6 × 2.4 × 2.0 (L×W×H) | 45.66 ± 3.90 | 72.2% | 43.2 | NA |
| **14** | one 1-layer trocar device (End-of-life) | LNG | 3 tablets 4.0 × 1.8 × 0.05 (L×W×H) | 13.6 × 2.4 × 2.0 (L×W×H) | 1.22[c] | 1.8% | 43.2 | NA |
| **15** | one 2-layer trocar device (sandwich) | LNG | 6 tablets 4.0 × 1.8 × 1.0 (L×W×H) | 13.6 × 2.4 × 2.1 (L×W×H) | 45.66 ± 3.90 | 68.8% | 86.4 | NA |
| **16** | one 2-layer trocar device (encapsulated) | LNG | 6 tablets 4.0 × 1.8 × 1.0 (L×W×H) | 13.6 × 2.4 × 2.1 (L×W×H) | 45.66 ± 3.90 | 68.8% | 86.4 | NA |

[a] LNG: levonorgestrel

[b] Dimensions provided in millimeters and specified as diameter and height (D, H) for cylindrical tablets and length, width and height (L, W, H) for rectangular tablets.

[c] Calculated drug loading determined by $Mass = Avg. LNG\ density\ [mg/mm^3] \times LNG\ volume\ [mm^3]$

**Table S2**. Detailed comparison of contraceptive MoSAIC designs with Jadelle® and Nexplanon®.

| | **Jadelle® (Bayer)** | **Nexplanon® (Merck)** | **MoSAIC (Trocar designs)** | **MoSAIC (18G designs)** |
|---|---|---|---|---|
| **Mechanism** | Diffusion | Diffusion | Surface area-controlled dissolution | Surface area-controlled dissolution |
| **Dimensions** | 43 mm length<br>2.5 mm diam.<br>2 rods | 40 mm length<br>2 mm diam.<br>1 rod | 27.9 mm length<br>2.0 mm height<br>2.4 mm width<br>1 or 2 rods | 40.4 mm length<br>0.64 mm height<br>0.84 mm width<br>2 or 3 rods |
| **Drug** | Levonorgestrel | Etonogestrel | Levonorgestrel | Levonorgestrel |
| **Duration** | 5 years | 3 years | ~5 years (1 rod)<br>~10 years (2 rods) | ~1 year (2 rods)<br>~1.5 years (3 rods) |
| **Potential Advantages** | • Approved product<br>• Retievable | • Approved product<br>• Retrievable | • Biodegradable (no need for surgical retrieval)<br>• Minimal tail<br>• Retrievable<br>• Smaller form factor | • Biodegradable (no need for surgical retrieval)<br>• Minimal tail<br>• Retrievable<br>• Smaller form factor<br>• Administration with standard 18G needle |
| **Disadvantages** | • Requires surgical removal<br>• Requires 2 rods for 5 years | • Requires surgical removal | • Pre-clinical early development | • Pre-clinical early development |

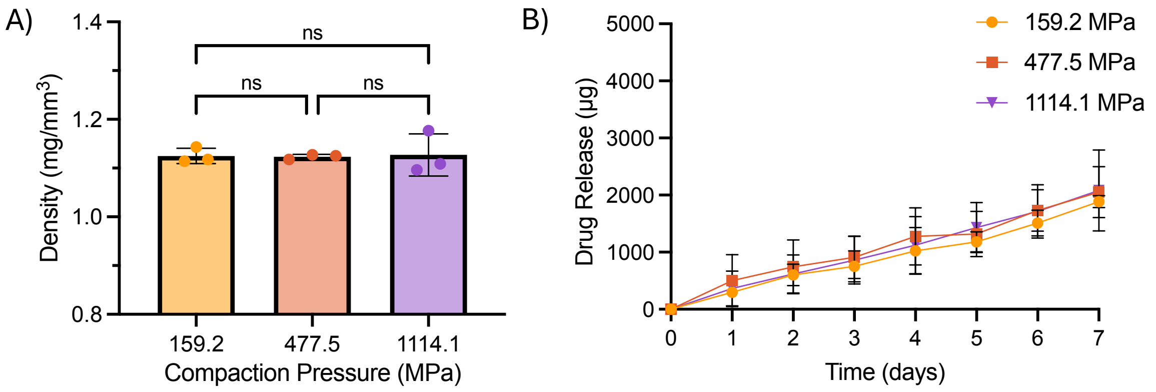


**Figure S1**. In vitro characterization of compacted LNG formulations. A) Effect of compaction force on the density of the solid LNG formulations. Formulations were 2.0 ± 0.01 mm in diameter, 1.26 ± 0.22 mm in height, and 4.57 ± 0.16 mg in weight. Compaction forces evaluated were 500 N, 1500 N, and 3500 N, which corresponds to compaction pressures of 159.16 MPa, 477.47 MPa, and 1114.11 MPa, respectively. Data presented as mean ± SD (n=3). B) In vitro drug release kinetics of compacted LNG formulations. The cylindrical formulations were incubated in 40 mL of pH 7.4 PBS supplemented with 10% 2-hydroxypropyl beta cyclodextrin (HP-β-CD) at 37°C with constant agitation. Drug concentrations in the samples were measured using HPLC. Data presented as mean ± SD (n=3).

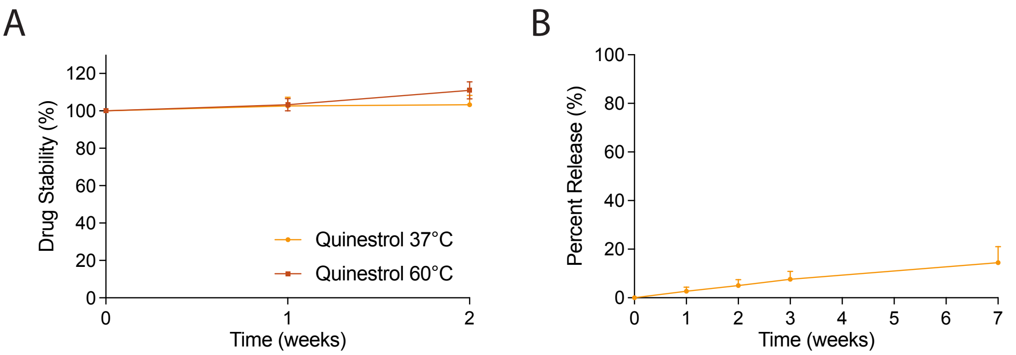


**Figure S2.** Quinestrol formulations. A) Quinestrol stability following incubation of solubilized drug at 37°C and 60°C for up to 2 weeks. Quinestrol was dissolved at a concentration of 0.551 ± 0.078 mg/mL in pH 7.4 PBS supplemented with 10% HP-β-CD. Data presented as mean ± SD (n=3). B) In vitro drug release kinetics of compacted LNG formulations. The cylindrical formulations weighing 26.73 ± 3.95 mg were incubated in 20 mL of pH 7.4 PBS supplemented with 10% HP-β-CD at 37°C with constant agitation. 10 mL of release medium was sampled at each timepoint and replaced with 10 mL of fresh medium. Drug concentrations in the samples were measured using HPLC. Data presented as mean ± SD. Data presented as mean ± SD (n=3).

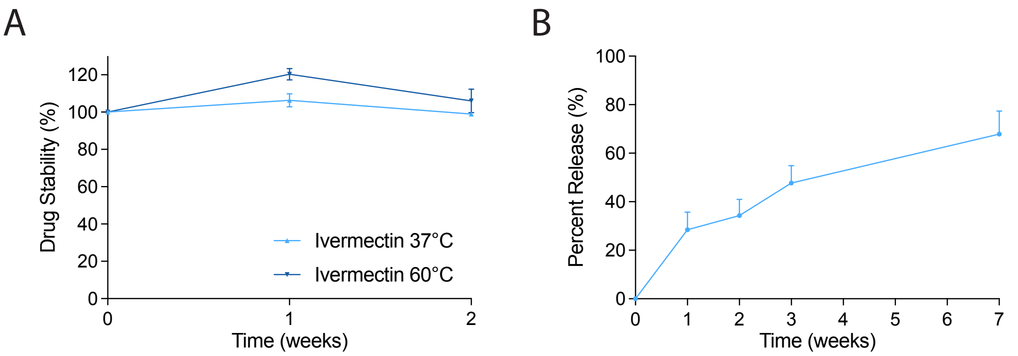


**Figure S3.** Ivermectin formulations. A) Ivermectin stability following incubation of solubilized drug at 37°C and 60°C for up to 2 weeks. Ivermectin was dissolved at a concentration of 1.047 ± 0.287 mg/mL in pH 7.4 PBS supplemented with 10% SDS. Data presented as mean ± SD (n=3). B) In vitro drug release kinetics of compacted LNG formulations. The cylindrical formulations weighing 28.22 ± 1.75 mg were incubated in 20 mL of pH 7.4 PBS supplemented with 10% SDS at 37°C with constant agitation. 10 mL of release medium was sampled at each timepoint and replaced with 10 mL of fresh medium. Drug concentrations in the samples were measured using HPLC. Data presented as mean ± SD. Data presented as mean ± SD (n=3).

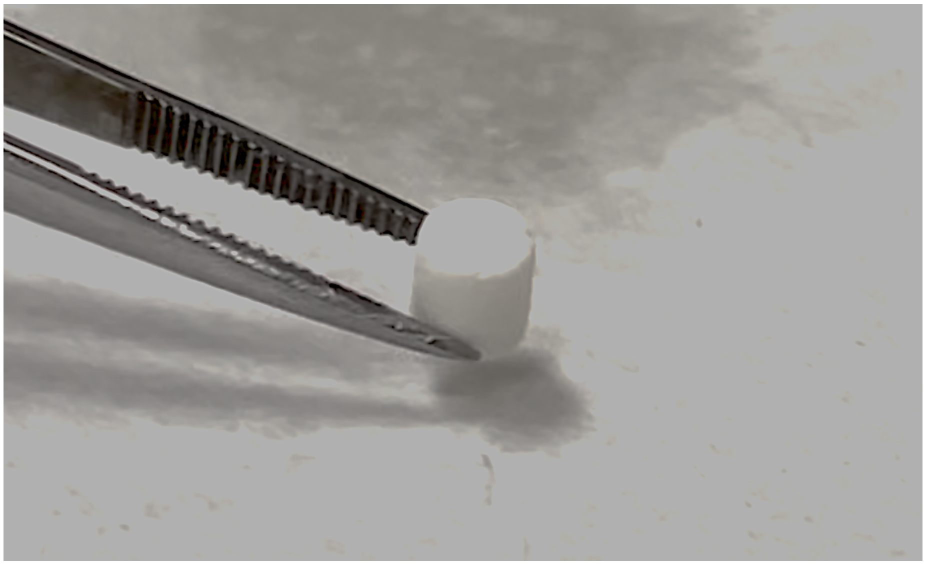

**Figure S4.** Photograph of manipulation of a retrieved compacted 4 mm x 4 mm cylindrical LNG formulation.

A

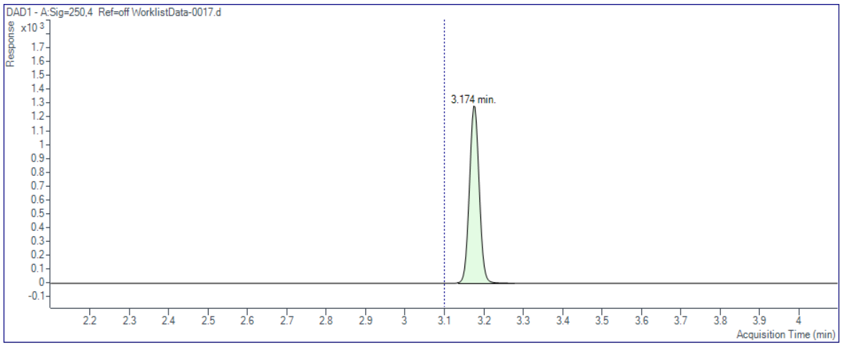


B

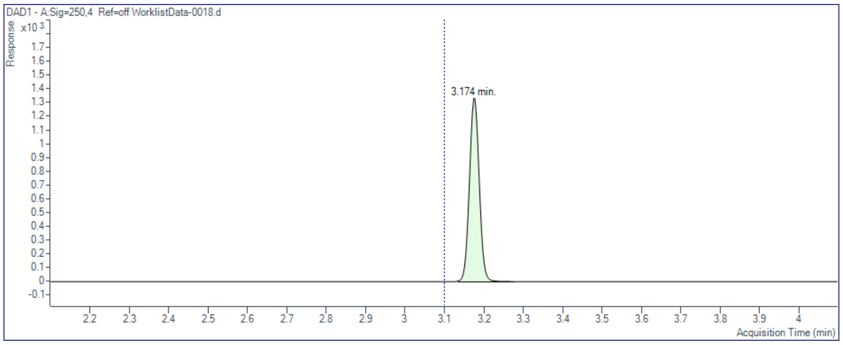


**Figure S5.** Representative HPLC chromatograms of solid LNG formulations before (A) and after (B) heating at 200°C. Retention time and purity remained consistent at 3.174 minutes and 100% respectively, suggesting that LNG remained stable following heat treatment.

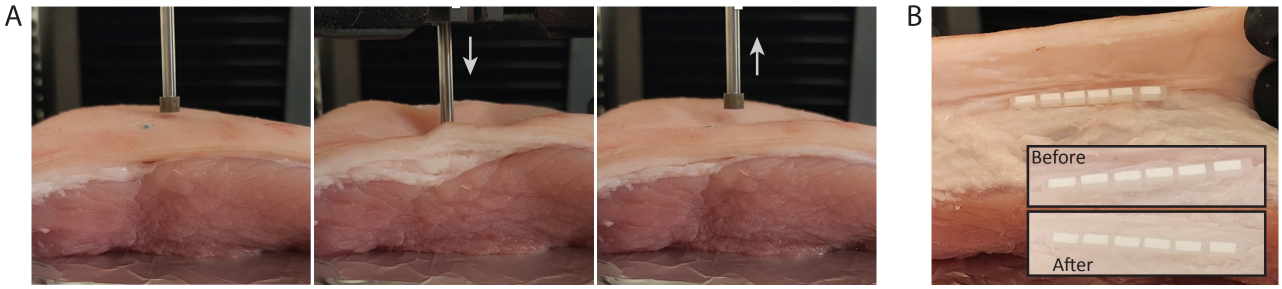


**Figure S6.** Photographs showing structural integrity of MoSAIC implants. A) The MoSAIC implant was inserted into a 1-inch-thick section of porcine skin tissue mounted on a rigid support. A 60 N point load was applied overtop of the center of the device using a 1/8" diameter steel plunger at a rate of 15 mm/min. B) Photograph of the intact MoSAIC device following force application.

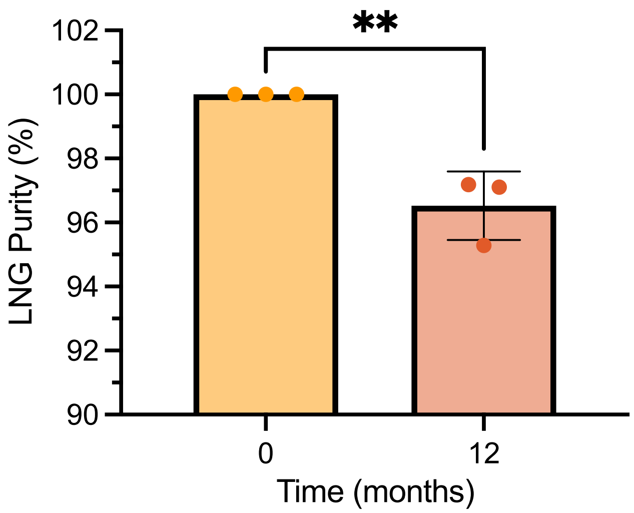


**Figure S7.** Chemical purity of LNG within monolithic implants that were implanted into the SC tissue of rats and retrieved after 1 year. Chemical purity is normalized to LNG within control monolithic implants that were not implanted into animals. Data presented as mean ± SD (n=3).

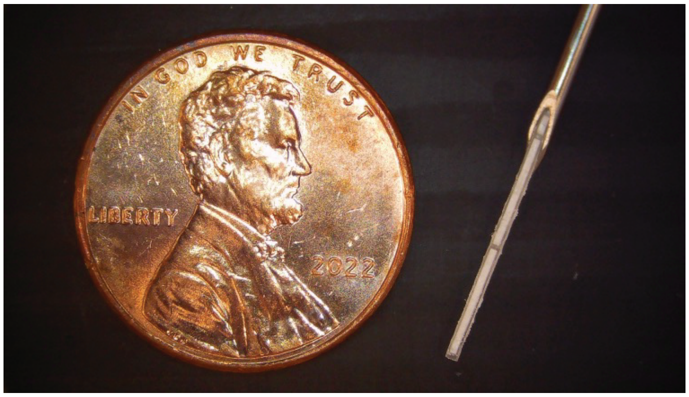


**Figure S8.** Photograph showing the concept of an 18G monolithic implant design. The implant is loaded with a compacted LNG formulation and fitted within the body of an 18-gauge hypodermic needle.
